# Non-muscle myosin-2 contractility-dependent actin turnover limits the length of epithelial microvilli

**DOI:** 10.1101/2020.05.01.072389

**Authors:** Colbie R. Chinowsky, Julia A. Pinette, Leslie M. Meenderink, Matthew J. Tyska

## Abstract

Epithelial brush borders are large arrays of microvilli that enable efficient solute uptake from luminal spaces. In the context of the intestinal tract, brush border microvilli drive functions that are critical for physiological homeostasis, including nutrient uptake and host defense. However, cytoskeletal mechanisms that regulate the assembly and morphology of these protrusions are poorly understood. The parallel actin bundles that support microvilli have their pointed-end rootlets anchored in a highly crosslinked filamentous meshwork referred to as the “terminal web”. Although classic EM studies revealed complex ultrastructure, the composition, organization, and function of the terminal web remains unclear. Here, we identify non-muscle myosin-2C (NM2C) as a major component of the brush border terminal web. NM2C is found in a dense, isotropic layer of puncta across the sub-apical domain, which transects the rootlets of microvillar actin bundles. Puncta in this network are separated by ∼210 nm, dimensions that are comparable to the expected size of filaments formed by NM2C. In primary intestinal organoid cultures, the terminal web NM2C network is highly dynamic and exhibits continuous remodeling. Using pharmacological and genetic perturbations to disrupt NM2C activity in cultured intestinal epithelial cells, we found that this motor controls the length of growing microvilli by regulating actin turnover in a manner that requires a fully active motor domain. Our findings answer a decades old question on the function of terminal web myosin and hold broad implications for understanding apical morphogenesis in diverse epithelial systems.

## INTRODUCTION

Hollow organs such as the intestinal track, kidney tubules, and brain ventricles, are lined with solute transporting epithelial cells. In the small intestine, individual epithelial cells, known as enterocytes, present thousands of microvilli on their apical surface. These protrusions are tightly packed into a highly ordered array collectively known as the brush border (1). Microvilli serve two general functions - they markedly increase the membrane surface area for nutrient absorption, and also provide the first line of defense against harmful compounds and microbes found in the luminal compartment (2,3). Due to the constant regeneration of the mammalian gut epithelium, continuous differentiation of enterocytes and assembly of the brush border are critical for maintaining homeostasis throughout an organism’s lifetime. Despite microvilli occupying a critical physiological interface, mechanisms that regulate apical morphogenesis remain unclear.

A single microvillus is supported by a core of 20-30 actin filaments, bundled together in parallel by actin bundlers villin, espin, and fimbrin (4–7), with the barbed-ends oriented at the distal tips, and the pointed-ends anchored in a dense, filamentous network known as the terminal web (8–10). While the terminal web was first described several decades ago in ultrastructural studies, little is known about how it contributes to the apical domain structure, microvillar organization, or brush border function. Interestingly, knockout (KO) mouse models lacking major brush border structural components, such as PACSIN-2, plastin-1, or actin bundling proteins (villin, espin and plastin-1) exhibit significant perturbations to the terminal web (11–13). All of these models exhibit a common phenotype, where the terminal web thins and microvilli become disorganized or shortened, which further suggests that terminal web architecture and brush border morphology are intimately linked.

Previous biochemical and ultrastructural studies of the terminal web identified several components of this filamentous network, including intermediate filaments and spectrins (14–17). Early deep-etch electron microscopy (EM) studies also identified “8 nm filaments”, which appeared to crosslink microvillar actin bundle rootlets (18). However, these structures did not label with myosin sub-fragment 1 (S1) suggesting they were not composed of actin. Additional immunogold labeling EM data led to speculation that 8 nm filaments contained myosin (9,19). This was further supported by immunofluorescence labeling studies that made use of a pan-myosin antibody raised against myosin-2 epitopes (20). However, the identity of the myosin, and the organization and function of 8 nm filaments in brush border structure and function remained unclear for decades.

All class 2 myosins are constitutive dimers of heavy chains, each comprised of an N-terminal motor domain, followed by a tandem pair of IQ motifs that are constitutively bound by regulatory and essential light chains (RLC and ELC, respectively), a long coiled-coil, and a non-helical tail piece at the C-terminus (21). A fully functional myosin-2 molecule consists of two heavy chains (HC), each bound by one RLC and one ELC. Class 2 myosins are regulated by reversible phosphorylation at sites on the RLC, the C-terminal tailpiece, or both, which controls activation and assembly of contractile units, respectively (22–24). Driven by electrostatic interactions between their coiled-coil tails, skeletal, cardiac and non-muscle myosin-2 (NM2) variants self-assemble into bipolar filaments that serve as fundamental force-generating units (25,26).

Non-muscle class 2 myosins (NM2A encoded by *MYH9*, NM2B encoded by *MYH10*, and NM2C encoded by *MYH14*) are ubiquitously expressed and have been implicated in diverse contractile activities including cytokinesis, cell motility, and the control of cell morphology (22). At higher levels of biological complexity, these motors have been implicated in shaping and bending of tissues, collective cell migration, and regulating the paracellular permeability of epithelial sheets (22,27). NM2 paralogs have distinct kinetic properties, which allow these motors to perform specific and distinct functions within the cell (28,29). In the context of *in vitro* sliding filament assays, tail-less NM2C heavy meromyosin (HMM) fragments move actin filaments at ∼0.05 µm/s, slower than both NM2B (∼0.08 µm/s) and NM2A (∼0.29 µm/s) (30). Bipolar filaments assembled by NM2C (∼293 nm) tend to be shorter than those assembled by NM2B (∼323 nm) or NM2A (∼301 nm). Filaments composed of NM2C also contain fewer molecules than those assembled by NM2A and NM2B, resulting in thinner filaments (∼8 nm vs. ∼11 nm wide) and presumably a lower potential for generating force (29).

A previous proteomic study by our laboratory revealed that brush border fractions isolated from mouse small intestine contain all three NM2 paralogs, although NM2C is by far the most abundant (31). NM2C remains the most poorly understood paralog with regard to biophysical properties and physiological function. NM2C exhibits specific expression to pituitary gland and glial cells, as well as inner ear sensory, intestinal, and kidney epithelia (32). Mutations in *MYH14* have been linked to hearing loss, peripheral neuropathies, and developmental defects in the lower gastrointestinal tract (33–41). The parallel perturbation of both inner ear and intestinal epithelial systems by mutations in *MYH14* is intriguing, as actin bundled-supported stereocilia found on the apical surface of hair cells are closely related to microvilli found on solute transporting epithelia, and may share mechanisms of assembly and maintenance (42,43). Indeed, previous studies of a NM2C-EGFP expressing mouse revealed that this motor is highly enriched at the junctional margins of sensory and intestinal epithelial cells, where cell-cell contacts are formed (44). At these sites, NM2 assembles into circumferential sarcomere-like structures, characterized by a linear array of puncta that are uniformly separated by ∼300-400 nm, a distance comparable to the expected length of contractile filaments formed by NM2 isoforms (29,44). Treatment of NM2C-EGFP mouse epithelial tissues with myosin-2 inhibitor blebbistatin (which prevents force generation) increased the inter-puncta distance, indicating that under normal conditions the circumferential NM2 band is under tension (44). Careful inspection of images in this previous study also revealed that NM2C signal is not confined to the circumferential junctional array, as dimmer puncta also appear to span the cell medially, throughout the entire sub-apical region.

In this paper, we report that NM2C localizes at the pointed ends of microvillar actin bundles, where it forms a highly dynamic array of puncta that are confined to the plane of the sub-apical terminal web. Although these puncta demonstrate ∼two-fold lower levels of NM2C enrichment relative to those found in circumferential array, based on their spacing and dynamics, we propose that they represent NM2C contractile filaments. Using pharmacological and genetic perturbations in cultured intestinal epithelial cells, we found that the activity of this motor controls the length of growing microvilli by regulating actin disassembly at the pointed ends. These findings address a decades old question on the function of the terminal web myosin-2 and hold important implications for understanding brush border assembly and apical morphogenesis in epithelial systems.

## RESULTS

### NM2C is enriched in the brush border terminal web *in vivo*

Previous studies established that NM2C localizes to peripheral cell-cell junctions in inner ear and intestinal tissues in mice (32,44). To investigate the sub-cellular distribution of NM2C in intestinal epithelial cells in more detail, we took advantage of mice that express an EGFP-tagged form of NM2C from the endogenous locus (44). In this model, EGFP is fused in frame to the C-terminus of the NM2C heavy chain. Confocal imaging of frozen intestinal tissue sections derived from these mice allowed us to visualize the strong apical signal of NM2C along the full length of villi (Figure 1A). Lower levels of NM2C were also observed on the apical surface of immature epithelial cells in the stem-cell containing crypt compartments (Figure 1A, merge). To gain additional resolution at the sub-cellular scale, tissue sections from NM2C-EGFP mice were subject to super-resolution structured illumination microscopy (SIM). SIM images of enterocytes were acquired with the apical-basolateral axis either parallel or perpendicular to the focal plane (lateral or *en face*, respectively). Lateral views revealed that the apical NM2C-EGFP signal observed in lower magnification images was confined to a band of signal that overlapped specifically with the rootlets of microvilli, near the pointed-ends of core actin bundles (Figure 1B,C). This band of signal spanned the entire apical surface at the level of the terminal web and its intensity increased significantly where it connected to the circumferential/junctional array at the margins of the cell (Figure 1C, white arrows). This was further confirmed with quantitative linescan analysis of NM2C-EGFP fluorescence intensities along individual microvilli visualized in SIM images (Figure 1D). *En face* images of NM2C-EGFP tissue samples revealed that terminal web NM2C signal was strikingly punctate; the differential enrichment of NM2C in medial vs. junctional puncta was also apparent in these images (Figure 1E). Intensity analysis revealed that medial puncta in the terminal web were ∼half the intensity of junctional puncta (Figure 1F). Moreover, nearest neighbor distance calculations showed that medial and junctional puncta are minimally separated by approximately the same distance, ∼210 nm (Figure 1G). Thus, in addition to its localization in the circumferential band at the margins of enterocytes, NM2C is also enriched throughout the terminal web, where it is well-positioned to interact directly with the rootlets of microvillar core actin bundles.

**Figure 1:**
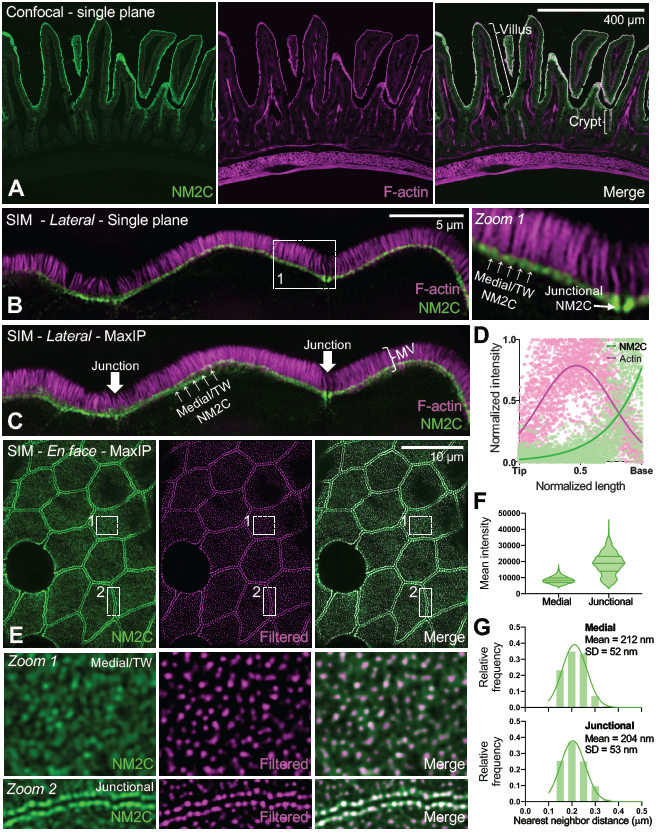
NM2C localizes to the enterocyte terminal web. **(A)** Single plane confocal microscopy of small intestinal tissue from a mouse endogenously expressing EGFP-NM2C (green), co-stained with phalliodin-568 to visualize F-actin (magenta). Representative villus and crypt regions are highlighted by brackets in merge panel. **(B)** Single plane structured illumination microscopy (SIM) image of small intestinal tissue from a mouse endogenously expressing EGFP-NM2C (green), co-stained with phalloidin-568 to visualize F-actin (magenta). Zoom 1 shows region bound by the dashed white box; medial and junctional population are highlighted. **(C)** Maximum intensity projection (MaxIP) of lateral view shown in B. **(D)** Normalized intensity plots of NM2C and F-signals taken from line-scans drawn parallel to the microvillar axis (n = 45) reveals enrichment of NM2C at the base of microvilli in the terminal web. **(E)** *En face* SIM MaxIP images of small intestinal tissue from a mouse endogenously expressing EGFP-NM2C (green); medial/terminal web and junctional populations are highlighted in zooms 1 and 2, respectively. SIM image (left) is shown in parallel with a version that was filtered using NanoF-SRRF (middle), which accentuated intensity peaks from individual NM2C puncta and allowed for more precise localization of their position. Merge image (right) shows composite of the original SIM image with the SRRF-filtered image. **(F)** Mean intensity of medial/terminal web vs. junctional NM2C puncta from SIM images. **(G)** Histograms of nearest neighbor distances generated by localizing medial (top) vs. junctional (bottom) NM2C puncta in SRRF-filtered SIM images. For F and G, n = 2,480 medial puncta and n = 1,019 junctional puncta. Scale is indicated on individual image panels.

### Terminal web and junctional NM2C puncta exhibit continuous remodeling

To enable live imaging studies of NM2C dynamics in the terminal web, we isolated crypts from NM2C-EGFP mice, which were then expanded into 2-dimensional (2D) organoid monolayers by plating on a thin layer of Matrigel. Confocal imaging of 2D organoids revealed a pattern of apical NM2C-EGFP distribution similar to that observed in fixed native tissue sections, with prominent junctional bands and a layer of medial puncta at the level of the terminal web (Figure 2A, Movie 1). Time-lapse imaging showed that both networks are highly dynamic and continuously remodeling, with puncta across the surface translocating, fusing and splitting on a timescale of minutes (Figure 2A zooms 1 and 2, Movie 1). We also performed photokinetic studies to examine the turnover rates of NM2C-EGFP puncta in the medial vs. junctional populations (Figure 2B-E). Fluorescence recovery after photobleaching (FRAP) analysis revealed that, despite the two-fold difference in puncta intensity (Figure 1F), the recovery rates for these two populations were remarkably similar (Figure 2D). Thus, medial NM2C-EGFP puncta exhibit spacing and turnover kinetics that are similar to junctional puncta. Given that the circumferential junctional belt of NM2 is an established contractile array (44), these data suggest that terminal web NM2C might also hold the potential to exert mechanical force.

**Figure 2:**
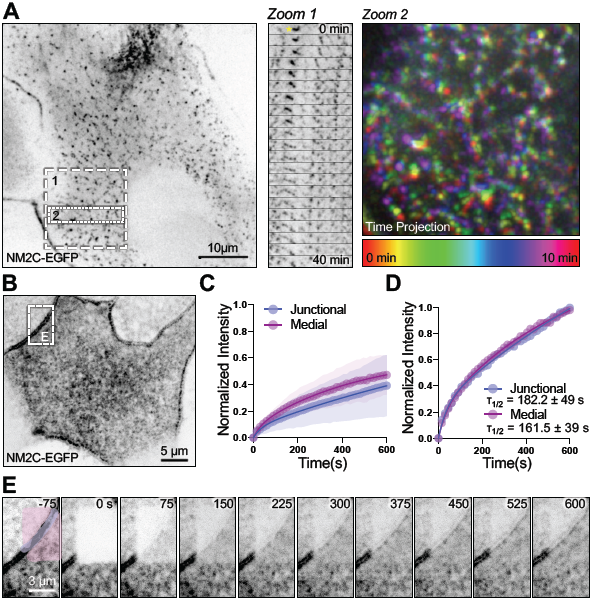
Medial and junctional populations NM2C puncta exhibit similar dynamics. **(A)** Live spinning disk confocal imaging of an organoid monolayer (i.e. 2D) derived from EGFP-NM2C expressing mouse small intestinal tissues; signal is inverted to facilitate visualization of dim structures. Zoom 1 shows a montage sampled at 2-minute intervals that reveals extensive remodeling of medial NM2C puncta over 40 minutes; puncta marked with yellow asterisk at t = 0 min exhibits striking expansion stretching/expansion during the time-lapse. Zoom 2 shows a color-coded time projection that reveals large-scale motion of medial NM2C network over 10 minutes. **(B)** FRAP analysis was performed on EGFP-NM2C puncta in 2D organoid monolayers to determine turnover rates in the junctional vs. medial populations. **(C)** FRAP recovery curves for junctional and medial NM2C populations normalized to pre-bleach intensity, show that both populations exhibit large immobile fractions (∼50%). **(D)** Kinetic analysis of datasets normalized to peak post-bleach intensities indicates that junctional and medial NM2C signal recovers at comparable rates; τ_1/2_ for both populations are shown on the plot. **(E)** Image montage of FRAP time-lapse data shown in B; junctional population is highlighted in purple, whereas medial NM2C is shown in pink.

### Activation of NM2 leads to shortening of microvilli

To further investigate the function of NM2C in the terminal web, we sought an appropriate cell culture model that recapitulates the terminal localization observed in native tissue. Based on previous studies from our laboratory and others (45,46), Ls174T-W4 (W4) cells mimic partially differentiated intestinal epithelial cell and thus, provide a model for studying actively growing microvilli (13,47). This cell line is derived from human colonic epithelium and can be induced via doxycycline treatment to acquire apical-basolateral polarity and assemble a brush border (46,48). In this model, microvilli extend from the cell surface parallel to the focal plane, which facilitates length measurements and assessment of protein localization along the protrusion axis. Of critical importance, W4 cells do not grow in monolayers and polarize as single cells. This unique advantage allowed us to specifically interrogate the function of the terminal web population of NM2C in the absence of the circumferential actin-myosin belt that normally assembles when cell-cell junctions are formed. SIM imaging of anti-NM2C stained W4 cells revealed striking localization at the base of microvilli, in a band that resembled the terminal web observed in native tissue samples (Supp. Figure 1A). Importantly, an NM2C construct with a C-terminal HaloTag also demonstrated similar terminal web enrichment in W4 cells (Supp. Figure 1B). Thus, both native and over-expressed NM2C localization in W4 cells are comparable to that observed in native intestinal tissue.

To probe NM2 function in the W4 cell terminal web, we first treated polarized W4 cells with calyculin A, a threonine/serine phosphatase inhibitor that is commonly used to elevate levels of phosphorylated myosin RLC and thus, increase active motor units (23,49–51). While calyculin A does not specifically target NM2C activity, our previous proteomic results (31,44) and antibody staining experiments in this study (Supp. Figure 1) indicate that this isoform is the dominant variant in the terminal web. Strikingly, treatment with 4 nM calyculin A resulted in marked shortening of microvilli over the course of 60 min (2.8 ± 0.6 µm control vs. 1.8 ± 0.5 µm calyculin A, Figure 3A-E, Movie 2). In calyculin A-treated W4 cells expressing Halo-NM2C, microvillar shortening was temporally paralleled by increased enrichment of NM2C in the terminal web (Figure 3D, Movie 2). In some cases, W4 cells appeared to lose nearly all surface microvilli following calyculin A treatment (Figure 3C). Indeed, the fraction of W4 cells that exhibited F-actin enriched brush borders also decreased significantly over the 60 min course of these experiments (Figure 3F). To further confirm that microvillar shortening was due to NM2 activation, we performed similar experiments using 4-hydroxyacetophenone (4-HAP), a small molecule that preferentially increases the activity of NM2B and NM2C, but not NM2A (52). In previous studies, 4-HAP was shown to increase NM2 motor activity and also enhance cortical enrichment of this motor (52,53). Similar to calyculin A-treated cells, W4 cells treated with 4-HAP exhibited increased terminal web enrichment of NM2C, and significant time-dependent shortening of microvilli and reduction in the percentage of cells presenting a brush border (Figure 3G-H, Movie 3). Together these drug treatment studies indicate that increased NM2 activity and enrichment in the terminal web is linked to microvillar shortening over time, and at longer time points, complete loss of the apical brush border.

**Figure 3:**
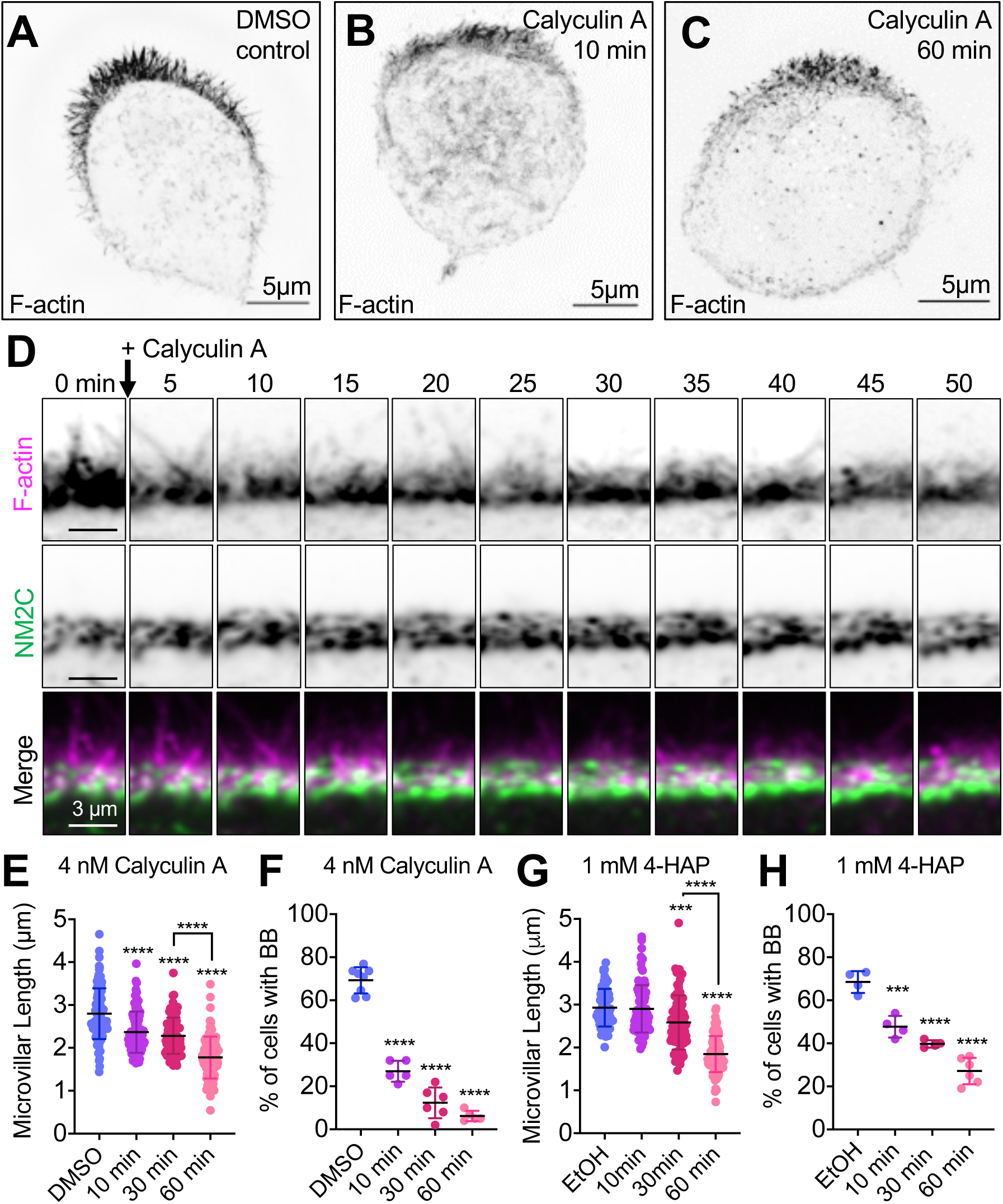
Activation of NM2 shortens epithelial microvilli. SIM MaxIP images of representative phalloidin-stained (F-actin) Ls174T-W4 cells fixed after **(A)** 60 min exposure to DMSO vehicle control, **(B)** 10 min exposure to 4 nM calyculin A, or **(C)** 60 min exposure to 4 nM calyculin A; signals are inverted to facilitate visualization of dim structures. **(D)** Image montage from spinning disk confocal time-lapse data shows the impact of calyculin A treatment on apical microvilli in a Ls174T-W4 cell expressing F-actin probe EGFP-UtrCH (magenta) with Halo-NM2C (green) labeled with JF585. **(E)** Quantification of microvillar length in Ls174T-W4 cells fixed after 10 min, 30 min, and 60 min in 4 nM calyculin A; each data point is a single microvillus, at least 10 microvilli per cell, minimum of 10 cells per condition. **(F)** Quantification of the percentage of brush border positive cells as a function of time in calyculin A. Each data point is percentage of cells with a brush border in a stitched 5×5 40x image. **(G)** Quantification of microvillar length in Ls174T-W4 cells fixed after 10 min, 30 min, and 60 min in 1 mM 4-HAP; each data point is a single microvillus, minimum of 10 cells per condition, at least 10 microvilli per cell. **(H)** Quantification of the percentage of brush border positive cells as a function of time in 4-HAP. Each data point is percentage of cells with a brush border in a stitched 5×5 40x image. E, F, G, and H were tested using ordinary one-way ANOVA with Dunnett’s multiple comparisons; ***, p < 0.0002; ****, p < 0.000 vs. DMSO/EtOH negative controls unless overwise noted.

### Inhibition of NM2 leads to microvillar elongation

To investigate the impact of NM2 loss-of-function on the brush border actin cytoskeleton, we treated W4 cells with the well characterized NM2 inhibitor, blebbistatin, which binds to and locks the NM2 motor domain in a non-force producing state (54,55). After treating polarized W4 cells with 20 µM blebbistatin, we noted that microvilli became significantly less dynamic (Movie 4) and subsequently exhibited a marked elongation, in some cases doubling their length (2.8 ± 0.6 µm control vs. 5.2 ± 1.3 µm blebbistatin treated) over the 60 min experimental time course (Figure 4A-E). In contrast the accumulation of NM2C observed in response to treatment with NM2 activators, inhibition with blebbistatin promoted a relatively static population of NM2C in the terminal web (Figure 4F, Movie 4). Moreover, although W4 cell microvilli normally extend from a single clearly defined apical ‘cap’, we noted that blebbistatin induced the dispersion of protrusions across the cell surface (Figure 4C). Thus, inhibition of NM2 leads to unregulated microvillar growth and disorganization of the brush border on the epithelial cell surface.

**Figure 4:**
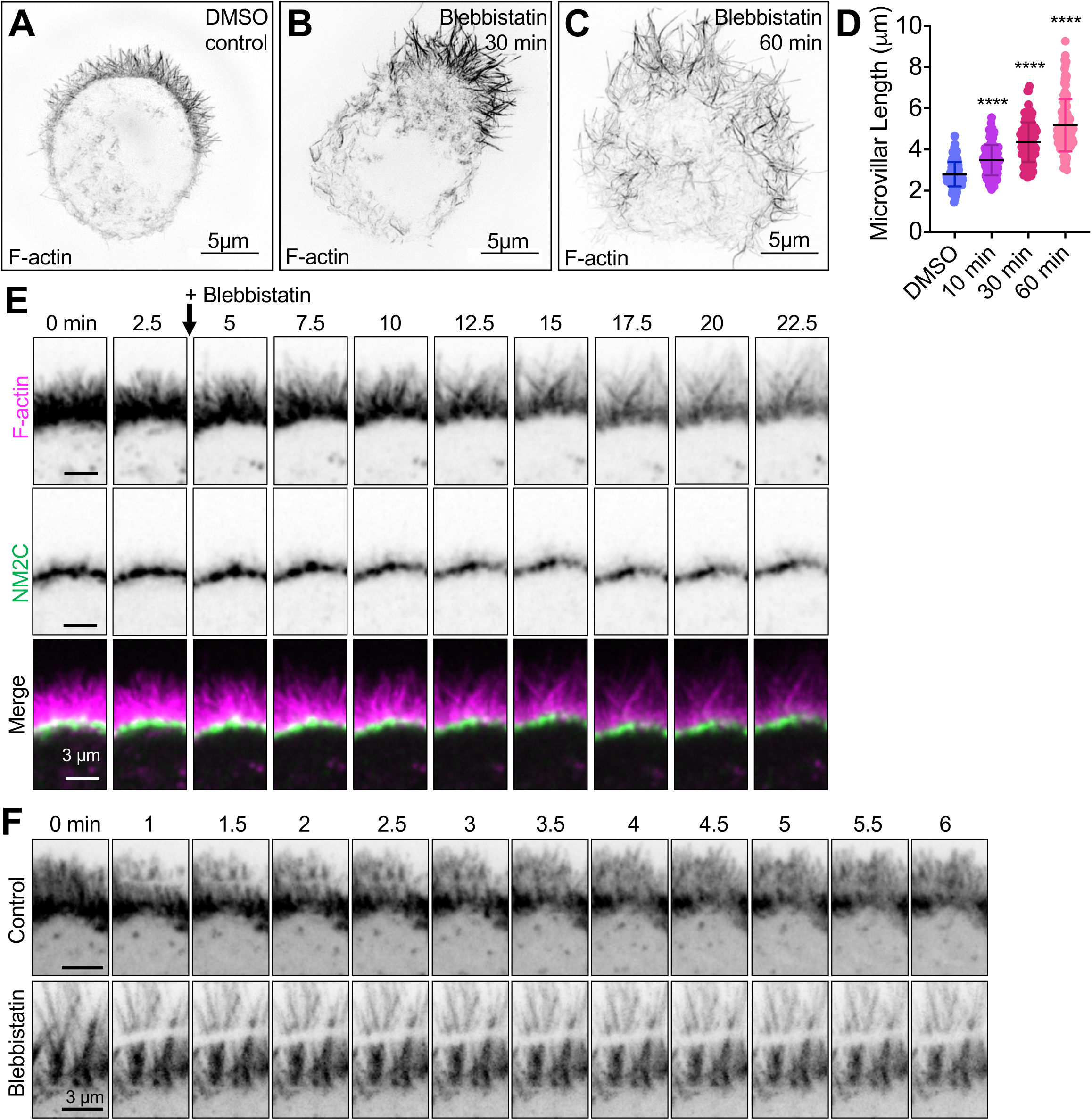
Inhibition of NM2 elongates microvilli and limits actin turnover. SIM MaxIP images of representative phalloidin-stained (F-actin) Ls174T-W4 cells fixed after **(A)** 60 min exposure to 20 μM DMSO vehicle control, **(B)** 10 min exposure to 20 μM blebbistatin, or **(C)** 60 min exposure to 20 μM blebbistatin; signals are inverted to facilitate visualization of dim structures. **(D)** Quantification of microvillar length from Ls174T-W4 cells fixed after 10 min, 30 min, and 60 min in 20 µM blebbistatin; each data point is a single microvillus, minimum of 10 cells per condition, at least 10 microvilli per cell. Ordinary one-way ANOVA with Dunnett’s multiple comparisons test; ****, p < 0.0001 vs. DMSO control. **(E)** Image montage of the brush border of a Ls174T-W4 cell expressing EGFP-UtrCH (magenta) and Halo-NM2C labeled with JF585 (green), imaged for 22.5 min after the addition of blebbistatin using spinning disk confocal microscopy; 2.5 min interval between frames. **(F)** Image montage of FRAP analysis of Ls174T-W4 cells expressing mNeon-Green β-actin in the absence (top row) or presence of 20 µM blebbistatin (bottom row) for 15 minutes prior to imaging.

### Blebbistatin reverses the impact of calyculin A on microvillar length

Blebbistatin is a well-established and specific inhibitor of NM2, whereas calyculin A has broader effects on a group of threonine/serine phosphatases (49,51,54,55). To determine if the microvillar shortening effects we observed with calyculin A could be countered with blebbistatin treatment, polarized W4 cells were subject to serial drug treatments (Movie 5). Cells were first treated with 4 nM calyculin A for 50 min until microvillar length was significantly diminished. At the 50 min mark, calyculin A was chased with 20 µM blebbistatin. These time-lapse data revealed that specific inhibition of NM2 with blebbistatin was sufficient to drive rapid elongation of the shortened microvilli that result from calyculin A treatment (Movie 5). Based on these data we conclude that the impact of calyculin A on microvillar length is most likely mediated by an increase in terminal web NM2 activity. In combination with the calyculin A and 4-HAP experiments described above, these data strongly suggest that under normal conditions, terminal web localized NM2C plays a role in limiting the length of microvilli and promoting their confinement in the apical domain.

### NM2 promotes actin turnover in brush border microvilli

How does terminal web NM2C limit the length of microvilli under normal conditions? Time-lapse analysis of W4 cells treated with 20 µM blebbistatin revealed that microvilli, which normally exhibit striking dynamics on the apical surface (e.g. elongation, shortening, and waving or pivoting around the base) immediately become static and subsequently begin to elongate (Movie 4). A previous study from our group established that the parallel actin bundles that support microvilli exhibit robust treadmilling (i.e. retrograde flow), where actin monomer incorporation at tip oriented barbed-ends is balanced by subunit removal from the pointed-ends that emerge from the base of these structures (56). Early in differentiation this treadmilling activity is coupled to directed motion of microvilli across the apical surface, which in turn promotes the adherent packing and organization of these protrusions (56). In treadmilling actin structures, steady-state length can be increased by raising the incorporation rate at the barbed-ends, lowering the disassembly rate at the pointed-ends, or some combination of the two (57–59). To examine the possibility that NM2C mechanical activity promotes the treadmilling of microvillar cores by driving the disassembly of actin bundles in the terminal web (where the pointed-ends reside), we performed FRAP on W4 cells transfected with mNeonGreen-β-actin, in the absence and presence of 20 µM blebbistatin. In control W4 cells, a rectangular ROI was bleached across the brush border and recovered rapidly, within < 4 minutes, presumably due to treadmilling of individual actin cores (Figure 4F, Movie 6). However, in W4 cells treated with 20 µM blebbistatin, bleached ROIs remained almost completely static and showed little to no recovery, even after up to 10 minutes (Figure 4F, Movie 7). Based on these findings, we conclude that under normal conditions, NM2C promotes the disassembly of the pointed-ends of core actin bundles within the terminal web. Without this activity (e.g. in response to blebbistatin treatment), treadmilling stalls and core bundles elongate as a result.

### NM2C motor domain activity is required for limiting microvillar length

All myosin motor domains contain highly conserved residues that participate directly in actin and ATP binding, as well as ATP hydrolysis and force production (21,27). Previous studies established that mutations to these residues disrupt myosin activity in predictable ways (60–64). To determine which facets of motor activity are needed for limiting microvillar length in W4 cells, we generated variants of NM2C with mutations predicted to lock the motor domain in weak or strong binding states (E497A or N252A, respectively)(60,62). We also targeted R784, which was identified as a critical residue in the NM2C crystal structure (65). R784 rests at the interface of the converter, N-terminal subdomain, and lever arm in the NM2C motor domain, and mutation of this residue leads to failure of converter rotation and impaired nucleotide binding, hydrolysis, and release (65). This general structure/function approach has been employed by our laboratory and others in the past to examine the requirement for myosin motor activity in different biological contexts (66–68). Typically, we would employ a KD/rescue strategy to first eliminate endogenous NM2C expression and then re-express NM2C mutant variants to assess their function. However, NM2C KD was not well tolerated by W4 cells (data not shown) as stable NM2C KD cells lines using lentiviral transduction of shRNAs were multinucleated and lacked a clearly defined apical brush border, which obscured analysis of microvillar morphology. As an alternate approach for assaying the impact of NM2C motor domain perturbations, we turned to an over-expression approach. Indeed, Halo-NM2C over-expression significantly shortened W4 cell microvilli (3.6 µm control vs 2.6 µm Halo-NM2C; Figure 5A,B,G). As a point of comparison, we also overexpressed EGFP-NM2A, which exerted an even more potent length reduction effect (3.6 µm control vs 2.3 µm in NM2A; Figure 5C,G). Strikingly, all three NM2C variants with function-disrupting mutations in the motor domain were unable to significantly reduce microvillar length when over-expressed in W4 cells (Figure 5D-G). From these data, we conclude that normal motor domain catalytic and mechanical activities are needed for NM2C to exert a length limiting effect on microvilli.

**Figure 5:**
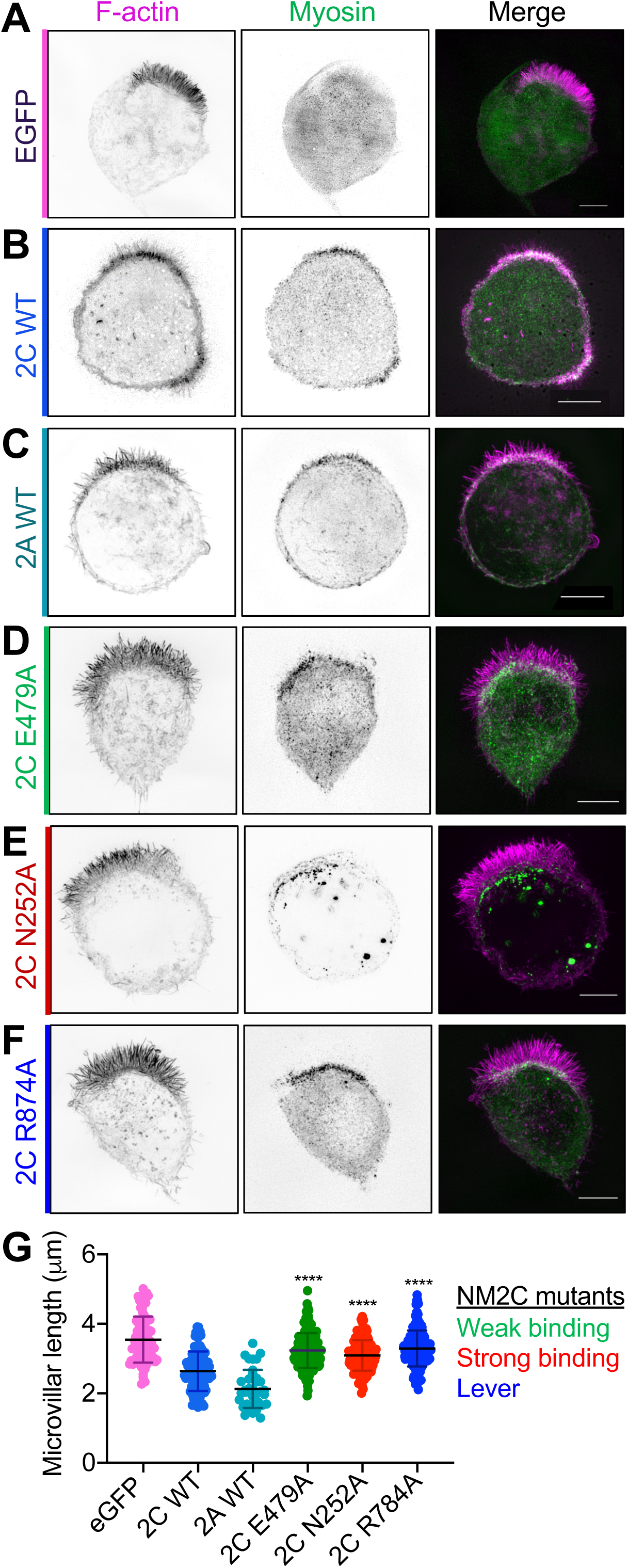
A functional NM2C motor domain is required for microvillar length regulation. SIM MaxIP images of representative Ls174T-W4 cell overexpressing **(A)** EGFP as a negative control, **(B)** WT Halo-NM2C, **(C)** WT EGFP-NM2A, **(D)** Halo-NM2C-E479A, **(E)** Halo-NM2C-N252A, or **(F)** Halo-NM2C-R874A. All Halo-NM2 constructs are labeled with JF585 (green) and cells are also co-stained with phalloidin to visualize F-actin (magenta). **(G)** Microvillar length quantification for each over-expression condition; measurements were made from 10 cells per construct, 8-10 microvilli measured per cell. Each data point is a single microvillus. Testing was performed using an ordinary one-way ANOVA with Dunnett’s multiple comparisons test; ****, p < 0.0001 vs. WT NM2C.

**Figure 6:**
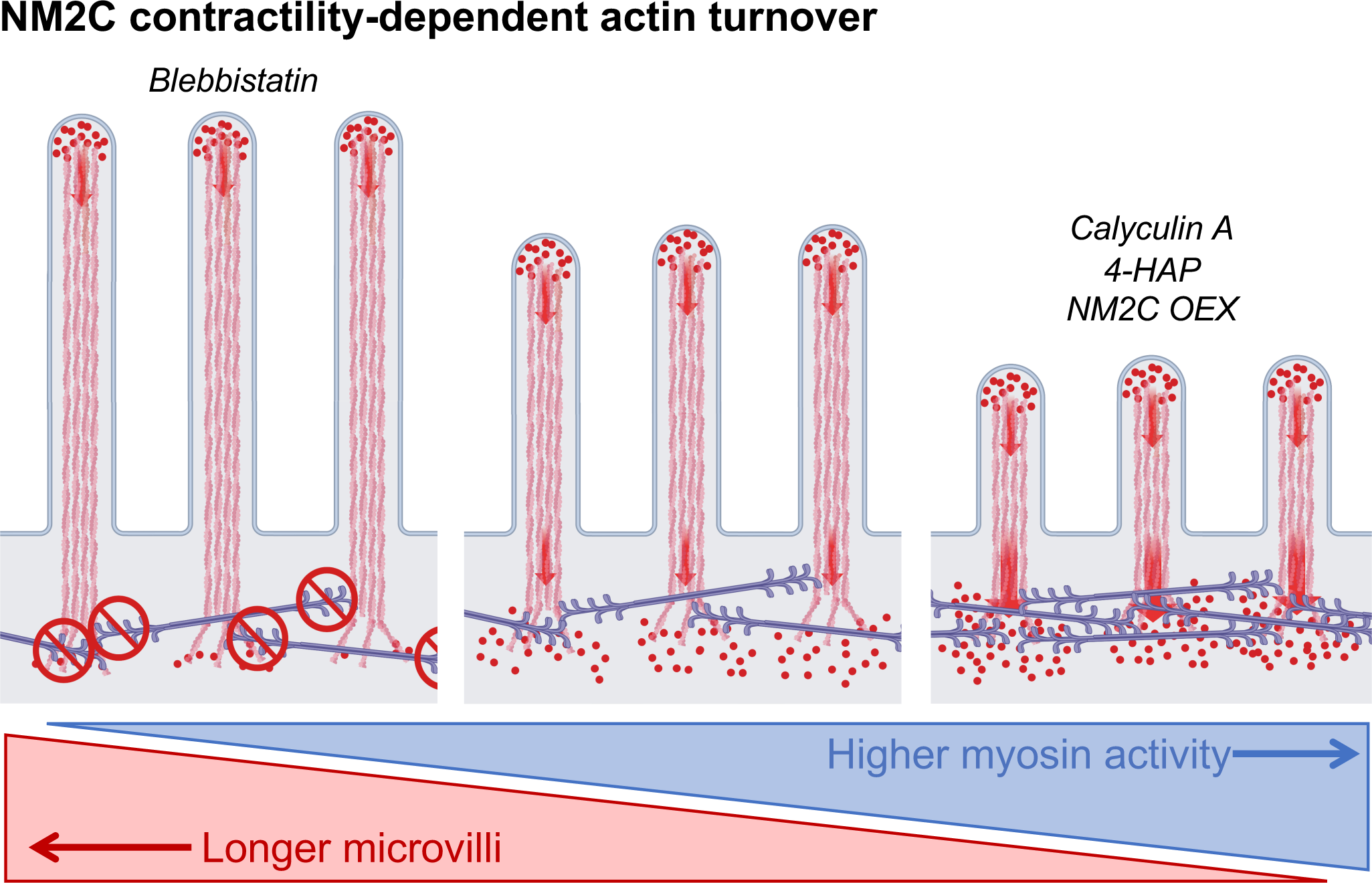
Model of contractility-dependent actin turnover in brush border microvilli. Cartoon summarizing the phenotyping observations linking NM2 inhibition, activation, and overexpression to perturbations to microvillus length. NM2C might also play a role in physically sequestering microvilli on the apical surface, a function that is not captured in this graphic summary.

## DISCUSSION

### NM2C activity limits the length of dynamic epithelial microvilli

In this work, we establish that the poorly studied NM2C isoform, which is highly expressed in transporting and sensory epithelia, not only localizes to the well-characterized junctional network, but is also enriched in a second network that spans the full diameter of the cell at the level of the sub-apical terminal web. Our super-resolution imaging studies show that terminal web NM2C is ideally positioned to interact directly with the rootlets of parallel actin bundles that support apical microvilli. Using a combination of chemical and genetic perturbations to increase or decrease the activity of this motor, we observed striking elongation or shortening of microvilli, respectively. We found the treadmilling dynamics, which are characteristic of nascent growing microvilli (56), are almost completely abolished upon treatment with the NM2 inhibitor, blebbistatin. Together these findings lead us to conclude that NM2C activity limits the length of microvilli, most likely by promoting disassembly of core bundle actin filaments near or at their pointed ends, which are embedded in the sub-apical terminal web. Because microvillar length control mechanisms are still poorly defined, these findings represent an important first step toward understanding how the dimensions of these structures are controlled.

### Organization of NM2C in the terminal web

The presence of a filament forming myosin in the terminal web was first suggested decades ago in high-resolution TEM images of microvillar rootlets (9). These iconic images revealed that the space between core bundle rootlets is spanned by small filaments with dimensions comparable to thick filaments found in muscle. Based on these early observations, Mooseker and Tilney postulated that adjacent microvilli were physically linked by a filament forming myosin, in a manner analogous to the organization of muscle sarcomeres (see Fig. 14 in ref 9). Indeed, the ability of NM2 to generate force at functionally significant levels is linked to its capacity to form bipolar filaments reminiscent of the thick filaments found in muscle sarcomeres (69–73). Based on *in vitro* studies with purified motor, NM2 filaments are composed of relatively small numbers (∼10s) of molecules, which self-associate via their long coiled-coil tail domains, leaving their N-terminal motor domains grouped at either end; the resulting structures extend 290-320 nm end-to-end (29). In this configuration, NM2 motor groups are capable of generating opposing forces of equal magnitude on actin structures bound by opposite ends of the bipolar filament, such as the rootlets of microvillar core bundles.

Our super-resolution imaging studies of native intestinal tissues revealed striking networks of NM2C puncta in both the circumferential junctional band (as previously described, 44), as well as the medial terminal web (Figure 1). Given that NM2C in the mouse model employed here is tagged on its C-terminus with EGFP (44), these puncta likely mark the center positions of bipolar filaments. Nearest neighbor measurements indicate that adjacent puncta in both junctional and medial populations are separated by comparable mean distances (204 vs. 212 nm, respectively; Figure 1), suggesting that these networks may be organized in similar ways. This is further supported by our FRAP studies of NM2C-EGFP dynamics in organoid monolayers, which showed that the turnover kinetics for these two populations is comparable. These similarities in spatial organization and turnover kinetics suggest that the junctional and medial populations of NM2C may be functionally analogous. Because the junctional band of NM2C is an established contractile array (44,74,75), these points might additionally argue that medial/terminal web NM2C is also capable of exerting force on the rootlets of microvillar actin core bundles. We also note here that the inter-puncta NM2C spacing measured in our samples (∼210 nm) is shorter than the mean length of NM2C filaments measured *in vitro* (∼290 nm), but comparable to the length of the NM2C filament bare zones (29). The super-resolution measurements of puncta spacing reported here are also smaller than previous measurements on the NM2C junctional network in intestinal or stomach epithelial tissues (∼400-500 nm)(44). The large range of spacing measurements from distinct biological contexts suggests a high degree of plasticity in NM2 contractile network organization, which probably reflects specific mechanical needs in these different environments.

### Contractility-dependent actin turnover as a conserved function for NM2

Our drug treatment studies with two activators (calyculin A and 4-HAP) and one inhibitor (blebbistatin) of NM2 activity demonstrate that tuning the level of myosin contractility in polarized epithelial cells has a profound impact on microvillar length, with higher activity leading to shorter protrusions. Of the three drug treatments applied in this study, only 4-HAP appears to exert some specificity toward NM2C; calyculin A and blebbistatin both exert effects on all NM2 isoforms (49,54). However, previous proteomic characterization of brush borders isolated from mouse small intestine indicate that NM2C is by far the most abundant isoform in this system (∼3-fold greater than NM2A, ∼12-fold greater than NM2B; 31). Moreover, our localization studies performed in W4 cells demonstrate that NM2C specifically enriches in the terminal web, whereas NM2A exhibits a non-polarized distribution over the entire cell cortex (Supp. Figure 1A vs. 1C). Taken together with our over-expression studies that establish NM2C is capable of shortening microvilli when present at high levels (Figure 5B,H), we conclude that the effects of drug treatments reported here primarily reflect an impact on NM2C activity in the terminal web.

The general concept that NM2 activity can promote actin network turnover is broadly consistent with findings from other diverse systems. Indeed, previous studies on mechanisms of neuronal growth cone motility showed that the rate of actin turnover (i.e. the treadmilling/retrograde flow rate) at the leading-edge decreases by ∼50% when cells are treated with blebbistatin (76). In this system, blebbistatin exposure also resulted in a striking elongation of filopodial actin bundles from their basal ends, which are normally embedded in a meshwork of lamellipodial actin filaments. NM2 contractility has also been implicated in the disassembly of actin filaments at the apical junctional complex in response to Ca^2+^ depletion (77), and in the recycling of filaments that is required for normal contractile ring constriction during cytokinesis (78,79). Thus, the NM2C-dependent microvillar length control mechanism we identify here represents a new facet of a broadly conserved class of function for filament-forming NM2 isoforms.

### Potential mechanisms for NM2-driven contractility-dependent actin turnover

How does NM2C activity promote the shortening of microvillar actin bundles? Previous work established that growing and nascent microvilli are supported by highly dynamic core bundles of parallel actin filaments, which collectively exhibit robust treadmilling (56,80). In treadmilling actin networks, new subunits incorporate at filament barbed-ends, whereas disassembly dominates at the pointed-ends. If assembly and disassembly rates are matched, steady-state length of the structure can remain constant. To shorten a treadmilling bundle, assembly rate must decrease, or disassembly rate must increase (or some combination of the two). Based on its localization, we propose that NM2C activity shortens microvilli by accelerating, either directly or indirectly, core actin bundle disassembly in the terminal web; under normal conditions this activity would contribute to actin bundle treadmilling and turnover. This is consistent with our live imaging observations showing that the robust turnover dynamics of W4 microvillar actin bundles is attenuated following blebbistatin treatment (Figure 4F).

NM2C might act directly on microvillar actin filament pointed-ends to drive their disassembly in the terminal web. Previous *in vitro* studies with purified actin and myosin-2 established that motor activity alone is capable of promoting filament disassembly (81). Interestingly, NM2C does exhibit kinetic properties (a moderate duty ratio and potentially strain-sensitive catalytic cycle) that would allow it to strain actin filaments (29). NM2C could also function indirectly by promoting the activity of other factors capable of disassembling filaments. This would be supported by recent studies on *Aplysia* growth cones which revealed that NM2 contractility can enhance the localization and severing activity of severing protein, cofilin (82). Proposed mechanisms by which NM2 might promote cofilin activity are based on its competition with cofilin binding, which creates distinct stiff and compliant zones (cofilin undecorated vs. decorated, respectively) along the filament, which are mechanically susceptible to severing at the boundaries of these regions (83–85). Although there is currently no data to support a role for cofilin in controlling the length of epithelial microvilli, this could be one area of focus for future studies.

### Additional roles for NM2C in polarized epithelia

In addition to promoting the turnover of actin filaments that comprise microvillar core bundles, NM2C might also participate in other roles critical for normal epithelial cell polarization and function. For example, we noticed that microvilli undergo a striking dispersion across the cell surface in response to blebbistatin treatment (Figure 4A-C, Movie 4). Thus, by binding directly to core actin bundles, NM2 might play a role in apicobasal polarity reinforcement and constraining of the size of the apical domain (86). NM2C could also contribute to the organelle exclusion properties of the terminal web. The high density of cytoskeletal (actin and intermediate) filaments in the terminal web creates a zone of organelle exclusion that prevents endomembrane compartments from making direct contact with the apical surface. If NM2C forms filaments that span the gaps between microvillar rootlets, these structures might participate in regulating the movement of vesicles through this region. Interestingly, the presence of a terminal web meshwork is a unique feature of the intestinal epithelium; other brush border presenting cells, such as those lining the kidney proximal tubule, lack a terminal web and zone of organelle exclusion (87,88). Importantly, these organs also lack strong NM2C localization, suggesting that NM2C may play a specific role in the compartmentalization of subcellular space in the intestinal epithelium (32).

Deeper insight on how NM2C contributes to epithelial physiology will come from careful analysis of *in vivo* loss-of-function models. Although a mouse model null for *MYH14* expression was previously described, the authors reported no overt physiological phenotypes (89). Other studies with the same KO mouse model revealed that in the absence of NM2C, NM2A and NM2B exhibit elevated levels in the intestinal epithelium as assessed by immunofluorescence signal (44). Thus, an apparent lack of overt phenotypes at the whole animal level may be due to functional compensation by NM2A and NM2B. Phenotypic characterization of the NM2C KO mouse represents an exciting direction for future studies, and might also provide a unique opportunity to investigate how the epithelium responds to higher levels of NM2 contractility, as would be expected based on the accelerated kinetic properties of NM2B and NM2A relative to NM2C (28,29).

## ACKNOWLEDGEMENTS

We thank all members of the Tyska laboratory, Vanderbilt Microtubule and Motors Club, and the Vanderbilt Epithelial Biology Center for advice and support. This work was supported in part by NIH grants R01-DK111949 and R01-DK095811 (PI: Tyska). Microscopy was performed in part through the Vanderbilt Cell Imaging Shared Resource, supported by the Vanderbilt Digestive Disease Research Center funded by NIH grant P30DK058404 (PI: Peek). CRC was supported in part by the Vanderbilt Molecular Biophysics Training Program T32-GM008320 (PI: Chazin).

## MATERIALS AND METHODS

### Frozen Tissue Preparation

Segments of small intestine where removed and flushed with cold PBS supplemented with 1.2 mM CaCl_2_ and 1 mM MgCl_2,_ and the fixed with 2% PFA/PBS for 15 minutes. After initial fixation, the intestinal tube was cut along its length and fixed for an additional 2 hrs in 2% PFA/PBS at room temperature with gentle agitation. Samples were washed with PBS three times and sub-dissected before being floated lumen-side down in 30% sucrose with 1% sodium azide/TBS at 4° C overnight. Samples were then streaked through OCT embedding medium before being oriented in a block filled with fresh OCT and snap-frozen in dry ice-cooled acetone. Samples were then sectioned (10 µm thick for confocal imaging or 2 µm thick for SIM imaging) and mounted on slides for phalloidin staining.

### Cell Culture

Ls174T-W4 (W4) cells (female *Hs* colon epithelial cells) were cultured in DMEM with high glucose and 2 mM L-glutamine supplemented with 10% tetracycline-free fetal bovine serum (FBS), blasticidin (10 µg/ml), G418 (1 mg/ml), and phleomycin (20 µg/ml); cells were grown at 37°C and 5% CO_2_ as our group and others have previously described. This cell line was generous gift from Dr. Hans Clevers (Utrecht University, Netherlands).

### Cloning and Constructs

pEGFP-NM2C was obtained from Addgene (plasmid #10843). mNEON-green-β-actin was purchased from Allele Biotechnology. pHalo-C1-NM2C (Halo-NM2C) construct was generated via PCR using pEGFP-NM2C as a template. NM2C open reading frame PCR product was TOPO cloned into the pCR8/GW/TOPO vector (K250020; Invitrogen) and then shuttled into the pHalo-C1 backbone, adapted for Gateway cloning using the Gateway conversion kit (11828029; Invitrogen). To generate pHalo-NM2C mutant variants, E479A, R257A, N252A, and R784A were introduced into pHalo-NM2C using QuikChange site-directed mutagenesis kit (200524; Agilent).

### Transfections

All transfections were performed using Lipofectamine2000 (#11668019; Invitrogen) according to manufacturer’s instructions and W4 cells were allowed to recover overnight. Following recovery, W4 cells were seeded onto plates or coverslips, and incubated overnight in the absence or presence of 1 μg/ml doxycycline to induce polarity, and then prepared for immunofluorescence or live cell imaging.

### Drug Treatments

For fix and stain experiments, W4 cells were split onto glass coverslips, and incubated for 12 hours in the presence of 1 μg/ml doxycycline to induce apicobasal polarity and brush border formation. Induced W4 cells were then incubated with either 20 µM Blebbistatin (B592500; Toronto Research Chemicals), 4 nM Calyculin A (PHZ1044; Invitrogen) or 1 mM 4-hydroxyacetophenone (278564; Sigma-Aldrich) for 10, 30 or 60 minutes. After drug incubation, cells were fixed using methods described below. For live cell imaging experiments, W4 cells were transfected with the appropriate construct and then split onto 35 mm glass bottom dishes (Invitro Scientific, D35-20– 1.5-N). Cells were imaged 24-72 hours after transfection. Once on the microscope, Blebbistatin, Calyculin A or 4-hydroxyacteophenone were added after ∼3-5 minutes of baseline imaging; cells were then imaged for an additional 20-60 minutes depending on the drug treatment.

### Cell and Tissue Staining

For SIM or laser scanning confocal imaging, W4 cells were plated onto glass coverslips, and allowed to adhere overnight in the presence of 1 μg/ml doxycycline. For NM2C staining, cells were then washed with warm phosphate-buffered saline (PBS) and fixed with warm 4% paraformaldehyde/PFA for 15 min at 37°C. Cells were subsequently washed three times with PBS, and then permeabilized with 0.1% Trition-X-100 in PBS for 15 minutes at room temperature. Cells were then blocked for 1 hr at room temperature in 5% bovine serum albumin (BSA) in PBS. Finally, cells were incubated with primary antibody (Proteintech anti-NM2C, #20716-1-AP) diluted in 5% BSA/PBS, at 37°C for 1.5 hr, followed by three washes with fresh PBS. Cells were then incubated for 1 hr with goat anti-rabbit Alexa Fluor 488F(ab’)2 Fragment (1:1000, A11070; Invitrogen) and Alexa Fluor 568 Phallodin (A12380; Invitrogen) at 37° C. Coverslips were washed three times with PBS and mounted on glass slides in ProLong Gold Antifade Mounting Media (P36930; Invitrogen). Frozen tissues sections were washed in PBS three times and stained with Alexa Fluor 568 Phallodin (A12380; Invitrogen) for 2 hrs at 37°C. Samples were then washed with PBS three times and mounted in ProLong Gold Antifade Mounting Media (P36930; Invitrogen). For cells transfected with Halo-tagged constructs, cells were incubated for 30 min with Janelia Fluor HaloTag ligand (gift from Dr. Luke Lavis, HHMI Janelia Farms) of the appropriate color, diluted to 100 nM after fixation with 4% PFA. Alexa Fluor 488-phalloidin (1:200, A12379; Invitrogen) was diluted in PBS and incubated for 1 hr at room temperature. Coverslips were washed three times with PBS then mounted on glass slides in ProLong Gold (P36930; Invitrogen).

### Light Microscopy and Image Processing

Super-resolution imaging was performed using a Nikon N-SIM Structured Illumination Microscope equipped with an Andor DU-897 EM-CCD, four excitation lasers (405, 488, 561, and 647 nm), and a 100x/1.49 NA TIRF objective. SIM images were reconstructed using Nikon Elements. For live-cell spinning disk confocal microscopy, W4 cells were transfected with the appropriate marker construct in a six well plate. Cells were then split onto 35 mm glass bottom dishes (Invitro Scientific, D35-20–1.5-N) and incubated in the presence or absence of 1 µg/ml doxycycline for a minimum of 12 hrs and up to 36 hrs. Cells transfected with Halo-tagged constructs were incubated with appropriately colored Janelia Fluor HaloTag ligand diluted to 100 nM in the standard cell culture media described above. Cells were incubated for 30 min with HaloTag ligand-media, media was then removed, and cells were washed with 37° C PBS. PBS was replaced with standard cell culture media and cells were allowed to equilibrate to new media for 15 min at 37° C. Live-cell imaging was performed on a Nikon Ti2 inverted light microscope equipped with a Yokogawa CSU-X1 spinning disk head, Andor DU-897 EMCCD camera or a Photometrics Prime 95B sCMOS camera, 488 nm and 561 nm excitation lasers, a 405 nm photo-stimulation laser directed by a Bruker mini-scanner to enable targeted photoactivation, photoconversion, and photobleaching), and a 100x/1.49 NA TIRF objective. Images were acquired every 15-60 sec for 20-60 min. For photokinetic studies of actin dynamics, baseline images were obtained for several frames prior to bleaching, then for an additional 3-10 min of recovery at 10 sec intervals. Bleaching was performed on a line ROI (0.1 μm in width and 3-15 μm in length) using a 405 nm laser at 30% power with a 10 μs dwell time. During imaging, cells were maintained with humidity at 37° C with 5% CO_2_ using a stage-top incubation system. Image acquisition was controlled with Nikon Elements software. 3D time series images were oversampled in the z-dimension with z-steps ranging from 0.09 μm to 0.18 μm to allow for deconvolution (Nikon Elements Automatic or Richardson-Lucy algorithms). Images were contrast enhanced, cropped, and aligned using ImageJ software (NIH) or Nikon Elements. Two-dimensional images were generally viewed as a maximum intensity projection (MaxIP); three-dimensional depth-coding was performed using ImageJ software (NIH).

### Quantification and Statistical Analysis

All image analysis was performed using FIJI or Nikon Elements software, and all quantitative data are from at least three independent experiments. To perform line-scan analysis, a line was drawn along the axis of microvilli that were in plane with a distinct tip and base visible. The intensity of NM2C signal was recorded and normalized with the lowest intensity set to 0 and the maximum set to 1. The microvillar length axis from individual scans was also normalized with the base set to 0 and the tip set to 1. Normalized line-scans were then plotted together and fit to a single Gaussian using nonlinear regression analysis (Prism v.7, GraphPad). For quantification of percentage of cells with brush border, cells were scored as ‘brush border positive’ if they displayed polarized F-actin accumulation as visualized using a 40X objective on a Nikon A1R laser-scanning confocal microscope as we have previously described. Microvillar length measurements were performed on projected SIM images by tracing individual microvillar actin bundles using FIJI. For analyses in which individual microvilli were measured, at least 10 microvillar actin bundles were scored per cell and at least 10 cells measured per experiment. Percent brush border and microvillar length data were analyzed with a D’Agostino and Pearson omnibus normality test to determine normal distribution. Normally distributed data were statistically analyzed to determine significance using the ordinary one-way ANOVA with Dunnett’s multiple comparisons to compare means between data sets. Statistical analyses performed are stated in the figure legends. All graphs were generated, and statistical analyses performed using Prism (v.7, GraphPad).

## Notes

### Competing Interest Statement

The authors have declared no competing interest.

## REFERENCES

1. Crawley SW, Mooseker MS, Tyska MJ. Shaping the intestinal brush border. J Cell Biol. 2014;207(4):441–51.

2. Helander HF, Fändriks L. Surface area of the digestive tract-revisited. Scand J Gastroenterol. 2014;49(6):681–9.

3. Vallance BA, Chan C, Robertson ML, Finlay BB. Enteropathogenic and enterohemorrhagic Escherichia coli infections: Emerging themes in pathogenesis and prevention. Can J Gastroenterol. 2002;16(11):771–8.

4. Bartles JR, Zheng L, Li A, Wierda A, Chen B. Small espin: A third actin-bundling protein and potential forked protein ortholog in brush border microvilli. J Cell Biol. 1998;143(1):107–19.

5. Mooseker MS, Graves TA, Wharton KA, Falco N, Howe CL. Regulation of Microvillus Structure : Calcium-dependent Solation and Cross-linking of Actin Filaments in the Microviili of Intestinal Epithelial Cells. J Cell Biol. 1980;(12).

6. Bretscher A, Weber K. Villin: The major microfilament-associated protein of the intestinal microvillus. Proc Natl Acad Sci U S A. 1979;76(5):2321–5.

7. Bretscher A, Weber K. Fimbrin, a new microfilament-associated protein present in microvilli and other cell surface structures. J Cell Biol. 1980;86(1):335–40.

8. Leblond CP, Puchtler H, Clermont Y. Structures corresponding to terminal bars and terminal web in many types of cells. Nature. 1960;186:784–8.

9. Mooseker MS, Tilney LG. Organization of an actin filament-membrane complex: Filament polarity and membrane attachment in the microvilli of intestinal epithelial cells. J Cell Biol. 1975;67(3):725–43.

10. Hull BE, Staehelin LA. The terminal web. A reevaluation of its structure and function. J Cell Biol. 1979;81(1):67–82.

11. Grimm-Gunter E-MS, Revenu C, Ramos S, Hurbain I, Smyth N, Ferrary E, et al. Plastin 1 Binds to Keratin and Is Required for Terminal Web Assembly in the Intestinal Epithelium. Mol Biol Cell. 2009;20:2673–83.

12. Revenu C, Ubelmann F, Hurbain I, El-Marjou F, Dingli F, Loew D, et al. A new role for the architecture of microvillar actin bundles in apical retention of membrane proteins. Mol Biol Cell. 2012;23:324–36.

13. Postema MM, Grega-Larson NE, Meenderink LM, Tyska MJ. PACSIN2-dependent apical endocytosis regulates the morphology of epithelial microvilli. Mol Biol Cell. 2019;30(19):2515–26.

14. Glenney JR, Glenney P. Fodrin is the general spectrin-like protein found in most cells whereas spectrin and the TW protein have a restricted distribution. Cell. 1983;

15. Glenney JR, Glenney P, Osborn M, Weber K. An F-actin- and calmodulin-binding protein from isolated intestinal brush borders has a morphology related to spectrin. Cell. 1982;28(4):843–54.

16. Mooseker MS, Bonder EM, Conzelman KA, Fishkind DJ, Howe CL, Keller TC. Brush border cytoskeleton and integration of cellular functions. J Cell Biol. 1984;99(1 II).

17. Howe CL, Sacramone LM, Mooseker MS, Morrow JS. Mechanisms of cytoskeletal regulation: Modulation of membrane affinity in avian brush border and erythrocyte spectrins. J Cell Biol. 1985;101(4):1379–85.

18. Hirokawa N, Heuser JE. Quick-freeze, deep-etch visualization of the cytoskeleton beneath surface differentiations of intestinal epithelial cells. J Cell Biol. 1981;91(2 Pt 1):399–409.

19. Hirokawa N, Tilney LG, Fujiwara K, Heuser JE. Organization of actin, myosin, and intermediate filaments in the brush border of intestinal epithelial cells. J Cell Biol. 1982;94(2):425–43.

20. Drenckhahn D, Gröschel-Stewart U. Localization of myosin, actin, and tropomyosin in rat intestinal epithelium: Immunohistochemical studies at the light and electron microscope levels. J Cell Biol. 1980;86(2):475–82.

21. Heissler SM, Manstein DJ. Nonmuscle myosin-2: Mix and match. Cell Mol Life Sci. 2013;70(1):1–21.

22. Betapudi V. Life without double-headed non-muscle myosin II motor proteins. Front Chem. 2014;2:1–13.

23. Dulyaninova NG, Bresnick AR. The heavy chain has its day: regulation of myosin-II assembly. Bioarchitecture. 2013;3(4):77–85.

24. Jana SS, Kim KY, Mao J, Kawamoto S, Sellers JR, Adelstein RS. An alternatively spliced isoform of non-muscle myosin II-C is not regulated by myosin light chain phosphorylation. J Biol Chem. 2009;284(17):11563–71.

25. McLachlan AD, Karn J. Periodic charge distributions in the myosin rod amino acid sequence match cross-bridge spacings in muscle. Nature. 1982;299(5880):226–31.

26. Atkinson SJ, Stewart M. Molecular interactions in myosin assembly. Role of the 28-residue charge repeat in the rod. J Mol Biol. 1992;226(1):7–13.

27. Chantler PD, Wylie SR, Wheeler-Jones CP, McGonnell IM. Conventional myosins - unconventional functions. Biophys Rev. 2010;2(2):67–82.

28. Heissler SM, Manstein DJ. Comparative kinetic and functional characterization of the motor domains of human nonmuscle myosin-2C isoforms. J Biol Chem. 2011;286(24):21191–202.

29. Billington N, Wang A, Mao J, Adelstein RS, Sellers JR. Characterization of three full-length human nonmuscle myosin II paralogs. J Biol Chem. 2013;288(46):33398–410.

30. Kim KY, Kovács M, Kawamoto S, Sellers JR, Adelstein RS. Disease-associated mutations and alternative splicing alter the enzymatic and motile activity of nonmuscle myosins II-B and II-C. J Biol Chem. 2005;280(24):22769–75.

31. McConnell RE, Benesh AE, Mao S, Tabb DL, Tyska MJ. Proteomic analysis of the enterocyte brush border. Am J Physiol - Gastrointest Liver Physiol. 2011;300(5):914–26.

32. Golomb E, Ma X, Jana SS, Preston YA, Kawamoto S, Shoham NG, et al. Identification and Characterization of Nonmuscle Myosin II-C, a New Member of the Myosin II Family. J Biol Chem. 2004;279(4):2800–8.

33. Kim SJ, Lee S, Park HJ, Kang TH, Sagong B, Baek JI, et al. Genetic association of MYH genes with hereditary hearing loss in Korea. Gene. 2016;592(1):177–82.

34. Han JJ, Nguyen PD, Oh DY, Han JH, Kim AR, Kim MY, et al. Elucidation of the unique mutation spectrum of severe hearing loss in a Vietnamese pediatric population. Sci Rep. 2019;9(1):1–9.

35. Song MH, Jung J, Rim JH, Choi HJ, Lee HJ, Noh B, et al. Genetic Inheritance of Late-Onset, Down-Sloping Hearing Loss and Its Implications for Auditory Rehabilitation. Ear Hear. 2020;41(1):114–24.

36. Lerat J, Magdelaine C, Roux AF, Darnaud L, Beauvais-Dzugan H, Naud S, et al. Hearing loss in inherited peripheral neuropathies: Molecular diagnosis by NGS in a French series. Mol Genet Genomic Med. 2019;7(9):1–12.

37. Fu X, Zhang L, Jin Y, Sun X, Zhang A, Wen Z, et al. Loss of Myh14 Increases Susceptibility to Noise-Induced Hearing Loss in CBA/CaJ Mice. Neural Plast. 2016;2016:1–16.

38. Zhu Z, Peng L, Chen G, Jiang W, Shen Z, Du C, et al. Mutations of MYH14 are associated to anorectal malformations with recto-perineal fistulas in a small subset of Chinese population. Clin Genet. 2017;92(5):503–9.

39. Donaudy F, Snoeckx R, Pfister M, Zenner HP, Blin N, Di Stazio M, et al. Nonmuscle Myosin Heavy-Chain Gene MYH14 Is Expressed in Cochlea and Mutated in Patients Affected by Autosomal Dominant Hearing Impairment (DFNA4). Am J Hum Genet. 2004;74(4):770–6.

40. Kim BJ, Kim AR, Han JH, Lee C, Oh DY, Choi BY. Discovery of MYH14 as an important and unique deafness gene causing prelingually severe autosomal dominant nonsyndromic hearing loss. J Gene Med. 2017;19(4):1–6.

41. Almutawa W, Smith C, Sabouny R, Smit RB, Zhao T, Wong R, et al. The R941L mutation in MYH14 disrupts mitochondrial fission and associates with peripheral neuropathy. EBioMedicine. 2019;45:379–92.

42. Mathur P, Yang J. Usher syndrome: Hearing loss, retinal degeneration and associated abnormalities. Biochim Biophys Acta - Mol Basis Dis. 2015;1852(3):406–20.

43. Prost J, Barbetta C, Joanny JF. Dynamical control of the shape and size of stereocilia and microvilli. Biophys J. 2007;93(4):1124–33.

44. Ebrahim S, Fujita T, Millis BA, Kozin E, Ma X, Baird MA, et al. NMII forms a contractile transcellular sarcomeric network to regulate apical cell junctions and tissue geometry. Curr Biol. 2014;23(8):731–6.

45. Grega-Larson NE, Crawley SW, Erwin AL, Tyska MJ. Cordon bleu promotes the assembly of brush border microvilli. Mol Biol Cell. 2015;26(21):3803–15.

46. Baas AF, Kuipers J, Wel NN Van Der, Batlle E, Koerten HK, Peters PJ, et al. Complete Polarization of Single Intestinal Epithelial Cells upon Activation of LKB1 by STRAD Annette. Cell. 2004;116:5.

47. Postema MM, Grega-Larson NE, Neininger AC, Tyska MJ. IRTKS (BAIAP2L1) Elongates Epithelial Microvilli Using EPS8-Dependent and Independent Mechanisms. Curr Biol. 2018;28(18):2876–88.

48. ten Klooster JP, Jansen M, Yuan J, Oorschot V, Begthel H, Di Giacomo V, et al. Mst4 and Ezrin Induce Brush Borders Downstream of the Lkb1/Strad/Mo25 Polarization Complex. Dev Cell. 2009;16(4):551–62.

49. Suzuki A, Itoh T. Effects of calyculin A on tension and myosin phosphorylation in skinned smooth muscle of the rabbit mesenteric artery. Br J Pharmacol. 1993;109(3):703–12.

50. Watanabe T, Hosoya H, Yonemura S. Regulation of Myosin II Dynamics by Phosphorylation and Dephosphorylation of Its Light Chain in Epithelial Cells. Mol Biol Cell. 2007;18(March):605–16.

51. Suganuma M, Fujiki H, Furuya-Suguri H, Yoshizawa S, Yasumoto S, Kato Y, et al. Calyculin A, an Inhibitor of Protein Phosphatases, a Potent Tumor Promoter on CD-I Mouse Skin. Cancer Res. 1990;50(12):3521–5.

52. Surcel A, Ng WP, West-Foyle H, Zhu Q, Ren Y, Avery LB, et al. Pharmacological activation of myosin II paralogs to correct cell mechanics defects. Proc Natl Acad Sci. 2015;112(5):1428–33.

53. Surcel A, Schiffhauer ES, Thomas DG, Zhu Q, DiNapoli KT, Herbig M, et al. Targeting mechanoresponsive proteins in pancreatic cancer: 4-hydroxyacetophenone blocks dissemination and invasion by activating MYH14. Cancer Res. 2019;79(18):4665–78.

54. Limouze J, Straight AF, Mitchison T, Sellers JR. Specificity of blebbistatin, an inhibitor of myosin II. J Muscle Res Cell Motil. 2004;25(4):337–41.

55. Kovács M, Tóth J, Hetényi C, Málnási-Csizmadia A, Seller JR. Mechanism of blebbistatin inhibition of myosin II. J Biol Chem. 2004;279(34):35557–63.

56. Meenderink LM, Gaeta IM, Postema MM, Cencer CS, Chinowsky CR, Krystofiak ES, et al. Actin Dynamics Drive Microvillar Motility and Clustering during Brush Border Assembly. Dev Cell. 2019;50(5).

57. Bonder EM, Fishkind DJ, Mooseker MS. Direct measurement of critical concentrations and assembly rate constants at the two ends of an actin filament. Cell. 1983;34(2):491–501.

58. Brenner SL, Korn ED. On the mechanism of actin monomer-polymer subunit exchange at steady state. J Biol Chem. 1983;258(8):5013–20.

59. Wang YL. Exchange of actin subunits at the leading edge of living fibroblasts: Possible role of treadmilling. J Cell Biol. 1985;101(2):597–602.

60. Sasaki N, Shimada T, Sutoh K. Mutational Analysis of the Switch II Loop of Dictyostelium Myosin II. J Biol Chem. 1998;273(32):20334–40.

61. Kambara T, Rhodes TE, Ikebe R, Yamada M, White HD, Ikebe M. Functional significance of the conserved residues in the flexible hinge region of the myosin motor domain. J Biol Chem. 1999;274(23):16400–6.

62. Shimada T, Sasaki N, Ohkura R, Sutoh K. Alanine scanning mutagenesis of the switch I region in the ATPase site of Dictyostelium discoideum myosin II. Biochemistry. 1997;36(46):14037–43.

63. Sasaki N, Ohkura R, Sutoh K. Dictyostelium myosin II mutations that uncouple the converter swing and ATP hydrolysis cycle. Biochemistry. 2003;42(1):90–5.

64. Kuhlman PA, Bagshaw CR. ATPase kinetics of the Dictyostelium discoideum myosin II motor domain. J Muscle Res Cell Motil. 1998;19(5):491–504.

65. Chinthalapudi K, Heissler SM, Preller M, Sellers JR, Manstein DJ. Mechanistic insights into the active site and allosteric communication pathways in human nonmuscle myosin-2C. Elife. 2017;6:e32742.

66. Weck ML, Crawley SW, Stone CR, Tyska MJ. Myosin-7b Promotes Distal Tip Localization of the Intermicrovillar Adhesion Complex. Curr Biol. 2016;26(20):2717–28.

67. Belyantseva IA, Boger ET, Naz S, Frolenkov GI, Sellers JR, Ahmed ZM, et al. Myosin-XVa is required for tip localization of whirlin and differential elongation of hair-cell stereocilia. Nat Cell Biol. 2005;7(2):148–56.

68. Sakai T, Umeki N, Ikebe R, Ikebe M. Cargo binding activates myosin VIIA motor function in cells. Proc Natl Acad Sci U S A. 2011;108(17):7028–33.

69. Stam S, Alberts J, Gardel ML, Munro E. Isoforms confer characteristic force generation and mechanosensation by myosin II filaments. Biophys J. 2015;108(8):1997–2006.

70. Huxley H. The Mechanism of Muscular Contraction. Science (80-). 1969;164(3886):1356–66.

71. Huxley H. The double array of filaments in cross-striated muscle. J Biophys Biochem Cytol. 1957 Sep;3(5):631–48.

72. Huxley H., Simmons R. Proposed Mechanism of Force Generation in Striated Muscle. Nature. 1971;233(5321):533–8.

73. Vicente-Manzanares M, Ma X, Adelstein RS, Horwitz AR. Non-muscle myosin II takes centre stage in cell adhesion and migration. Nat Rev Mol cell Biol. 2009;10(11):778–90.

74. Keller TCS, Conzelman KA, Chasan R, Mooseker MS. Role of myosin in terminal web contraction in isolated intestinal epithelial brush borders. J Cell Biol. 1985;100(5):1647–55.

75. Turner JR, Rill BK, Carlson SL, Carnes D, Kerner R, Mrsny RJ, et al. Physiological regulation of epithelial tight junctions is associated with myosin light-chain phosphorylation. Am J Physiol Physiol. 1997;273(4):C1378–85.

76. Medeiros NA, Burnette DT, Forscher P. Myosin II functions in actin-bundle turnover in neuronal growth cones. Nat Cell Biol. 2006;8(3):216–26.

77. Ivanov AI, McCall IC, Parkos CA, Nusrat A. Role for Actin Filament Turnover and a Myosin II Motor in Cytoskeleton-driven Disassembly of the Epithelial Apical Junctional Complex. Mol Biol Cell. 2004;15(6):2639–51.

78. Guha M, Zhou M, Wang Y. Cortical Actin Turnover during Cytokinesis Requires Myosin II. Curr Biol. 2005;15(8):732–6.

79. Murthy K, Wadsworth P. Myosin-II-Dependent Localization and Dynamics of F-Actin during Cytokinesis. Curr Biol. 2005;15(8):724–31.

80. Tyska MJ, Mooseker MS. MYO1A (Brush Border Myosin I) Dynamics in the Brush Border of LLC-PK1-CL4 Cells. Biophys J. 2002;82(4):1869–83.

81. Haviv L, Gillo D, Backouche F, Bernheim-Groswasser A. A Cytoskeletal Demolition Worker: Myosin II Acts as an Actin Depolymerization Agent. J Mol Biol. 2008;375(2):325–30.

82. Zhang XF, Ajeti V, Tsai N, Fereydooni A, Burns W, Murrell M, et al. Regulation of axon growth by myosin II–dependent mechanocatalysis of cofilin activity. J Cell Biol. 2019;218(7):2329–49.

83. De La Cruz EM, Martiel JL, Blanchoin L. Mechanical heterogeneity favors fragmentation of strained actin filaments. Biophys J. 2015;108(9):2270–81.

84. Schramm AC, Hocky GM, Voth GA, Blanchoin L, Martiel J-L, De La Cruz EM. Actin Filament Strain Promotes Severing and Cofilin Dissociation. Biophys J. 2017;112(12):2624–33.

85. R.McCullough B, Blanchoin L, Martiel J-L, De La Cruz EM. Cofilin Increases the Bending Flexibility of Actin Filaments: Implications for Severing and Cell Mechanics. J Mol Biol. 2008;381(3):550–8.

86. Ivanov AI, Hunt D, Utech M, Nusrat A, Parkos CA. Differential roles for actin polymerization and a myosin II motor in assembly of the epithelial apical junctional complex. Mol Biol Cell. 2005;16(6):2636–50.

87. Mooseker MS, Bonder EM, Conzelman KA, Fishkind DJ, Howe CL, Keller TC. The 0 cytoskeletal apparatus of the intestinal brush border. Ann Rev Cell Biol. 1985;1:209–41.

88. Coudrier E, Kerjaschki D, Louvard D. Cytoskeleton organization and submembranous interactions in intestinal and renal bursh borders. Kidney Int. 1988;34(3):309–20.

89. Ma X, Jana SS, Conti MA, Kawamoto S, Claycomb WC, Adelstein RS. Ablation of Nonmuscle Myosin II-B and II-C Reveals a Role for Nonmuscle Myosin II in Cardiac Myocyte Karyokinesis. Mol Biol Cell. 2010;21(24):4953–62.

