## Supplementary material for "Non-muscle myosin-2 contractility-dependent actin turnover limits the length of epithelial microvilli": Supp. Figures

### SUPPLEMENTAL FIGURE LEGENDS

**Supp. Figure 1 (related to Figure 3): NM2C localizes specifically to the terminal web of W4 cells.** SIM MaxIP of representative phalloidin-stained (F-actin, magenta) Ls175T-W4 cells displaying: **(A)** endogenous NM2C staining, **(B)** overexpression of Halo-NM2C labeled with JF585, or **(C)** overexpression of EGFP-NM2A. Scale bars are 5 µm.

**Supp. Figure 2 (Related to Figure 3): Activation of NM2C with 4-HAP shortens epithelial microvilli.** SIM MaxIP images of representative phalloidin-stained (F-actin) Ls174T-W4 cells fixed after: **(A)** 60 min exposure to EtOH vehicle control, **(B)** 10 min exposure to 1 mM 4-HAP, or **(C)** 60 min exposure to 1 mM 4-HAP; signals are inverted to facilitate visualization of dim structures. **(D)** Image montage from spinning disk confocal time-lapse data shows the impact of 4-HAP treatment on apical microvilli in a Ls174T-W4 cell expressing F-actin probe EGFP-UtrCH (magenta) with Halo-NM2C (green) labeled with JF585.

Supp. Figure 1 (related to Figure 3)

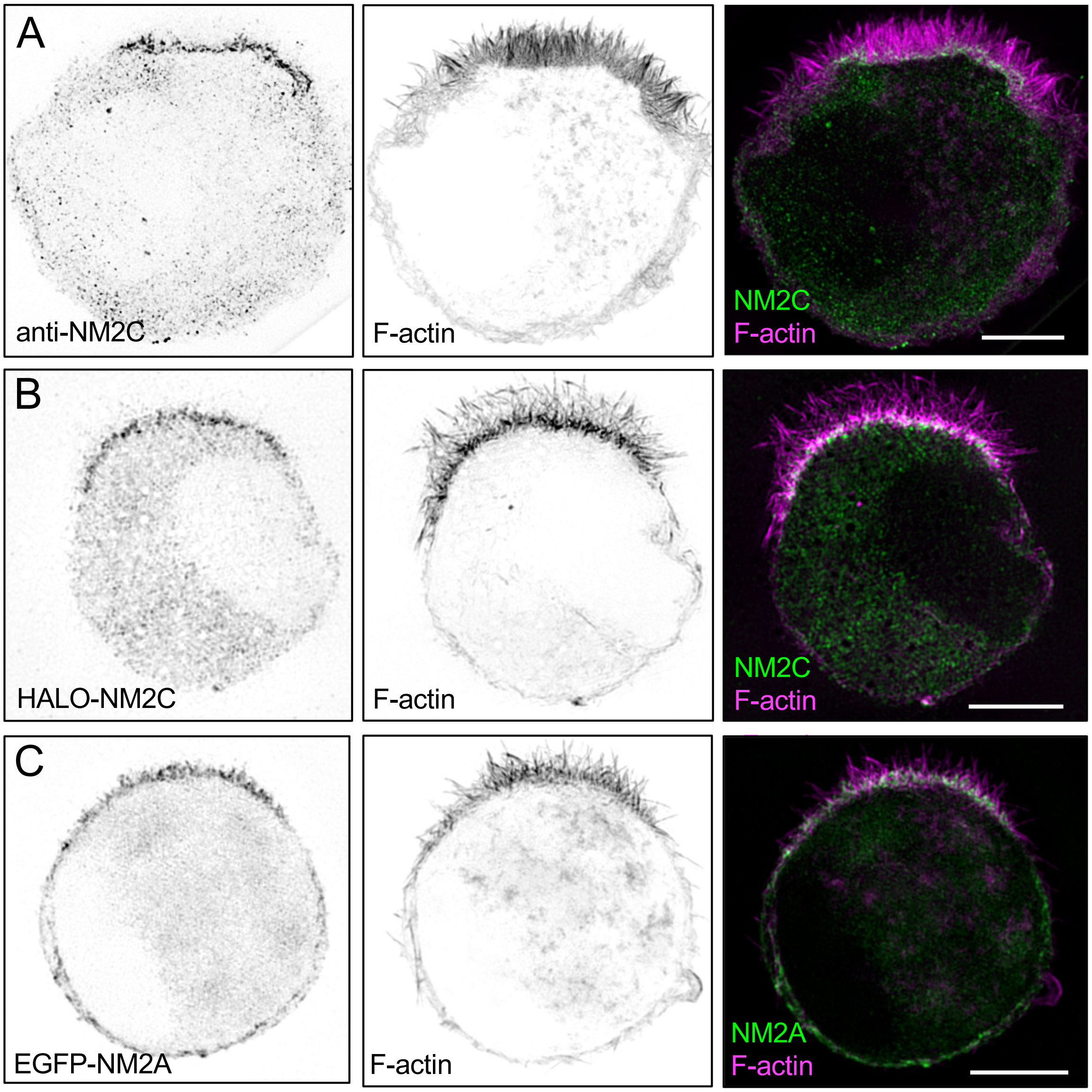

Supp. Figure 2 (Related to Figure 3)

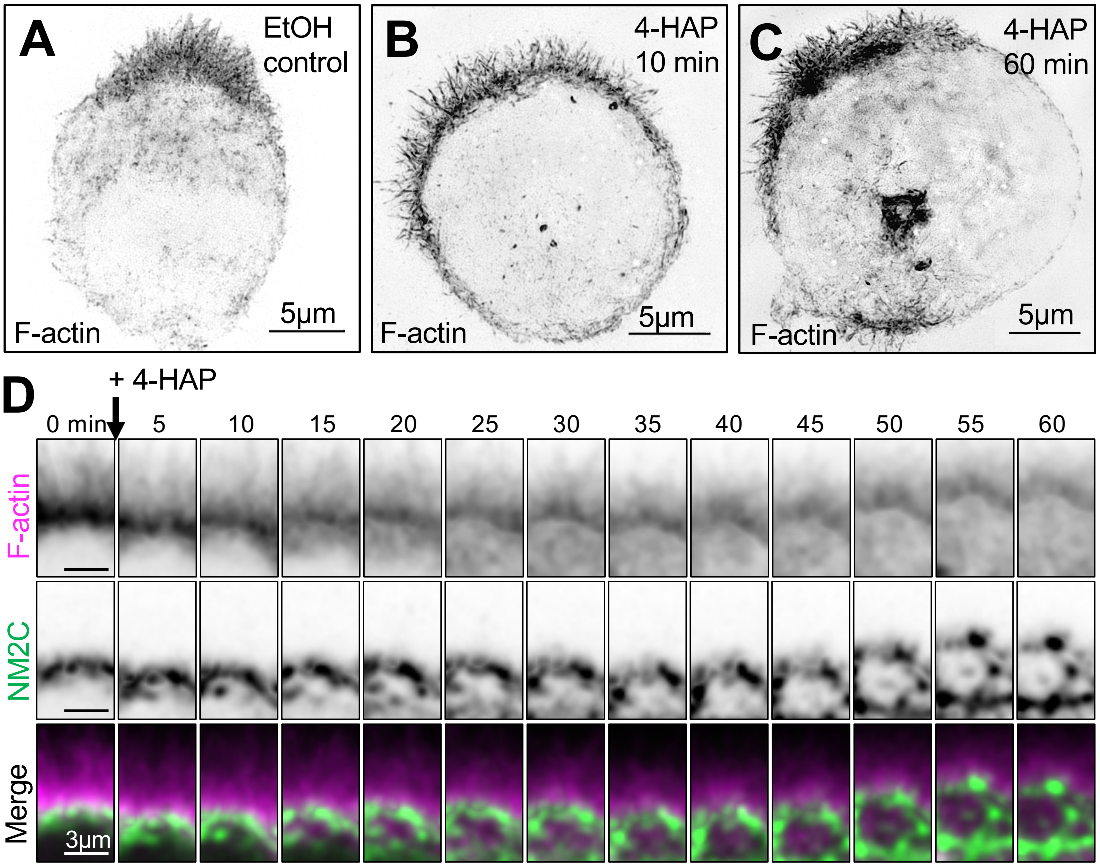
