## Supplementary material for "Non-muscle myosin-2 contractility-dependent actin turnover limits the length of epithelial microvilli": Movie Legends

**Movie 1 - Related to Figure 2. Live imaging of the apical surface of a 2D intestinal organoid monolayer derived from a NM2C-EGFP expressing mouse.** Spinning disk confocal images were acquired every 60 seconds for 40 minutes; playback is 15 frames per second; field width is 53 μm.

**Movie 2 - Related to Figure 3. Calyculin A shortens Ls174T-W4 cell microvilli.** Spinning disk confocal imaging of an induced Ls174T-W4 cell expressing EGFP-UtrCH (magenta) and Halo-NM2C labeled with JF585 (green). Images were acquired every 15 seconds for 60 minutes; playback is 15 frames per second; field width is 23 μm.

**Movie 3 - Related to Figure 3. 4-HAP shortens Ls174T-W4 cell microvilli.** Spinning disk confocal imaging of an induced Ls174T-W4 cell expressing EGFP-UtrCH (magenta) and Halo-NM2C labeled with JF585 (green). Movie was acquired every 15 seconds for 60 minutes; playback is 15 frames per second; field width is 32 μm.

**Movie 4 - Related to Figure 4. Blebbistatin elongates Ls174T-W4 cell microvilli.** Spinning disk confocal imaging of an induced Ls174T-W4 cell expressing EGFP-UtrCH (magenta) and Halo-NM2C labeled with JF585 (green). Images were acquired every 30 seconds for 30 minutes; playback is 5 frames per second; field width is 29 μm.

**Movie 5 - Blebbistatin rescues microvilli shortened by Calyculin A.** Spinning disk confocal imaging of an induced Ls174T-W4 cell expressing EGFP-UtrCH. Images were acquired every 30 seconds for 78 minutes; playback is 20 frames per second.

**Movie 6 - Related to Figure 4. Microvillar core actin bundles are rapidly turned over.** Spinning disk confocal imaging of a FRAP control experiment on induced Ls174T-W4 cell expressing mNeonGreen β-actin.  ROI is bleached using a 405-laser line at 30% power with 100 μs dwell time. 5 frames acquired pre-bleach at 60 second intervals;22 frames acquired post-bleach at 30 second intervals for 10 minutes. Playback is 3 frames per second.

**Movie 7 - Related to Figure 4. Inhibition of NM2 limits actin turnover.** Spinning disk confocal imaging of a FRAP experiment on induced Ls174T-W4 cell expressing mNeonGreen-β-actin, treated with blebbistatin for 15 minutes prior to imaging.  ROI is bleached using a 405-laser line at 30% power with 100 μs dwell time. 5 frames acquired pre-bleach at 60 second intervals; 22 frames acquired post-bleach at 30 second intervals for 10 minutes. Playback is 3 frames per second.
